# Multifractal Fluctuations in Electrogram Dynamics Distinguish Atrial Fibrillation Phenotype, Drug Response, and Imminent Termination: Implications for Mechanism and Treatment

**DOI:** 10.64898/2026.04.12.718068

**Authors:** Darius Chapman, Campbell Strong, Ivaylo Tonchev, Scott Lorensini, Sobhan Salari Shahrbabaki, Anand Ganesan

## Abstract

**Background:** Atrial fibrillation (AF) is maintained by complex dynamics, clinically characterised by bursting periods of organization and disorganization in intracardiac electrograms. We have previously postulated that cardiac conduction behaves like a critical system, where phase shift from organised rhythm to AF is a phase transition at the critical point. We thus hypothesized that using multifractal analysis of AF electrograms could potentially quantify non-stationary fluctuations, revealing novel mechanistic insights into the cardiac critical system and examine potential clinically relevant markers of AF dynamics, phenotype and treatment response.

**Objectives:** To determine whether multifractal analysis of AF electrograms can (i) Distinguish paroxysmal (PAF0 and non-paroxysmal AF (NPAF), (ii) predict response to pharmacologic modulation, and (iii) identify imminent spontaneous termination, thereby acting as marker of proximity to criticality along complex system phase spectrum.

**Methods:** We analysed >1.4 million seconds of high-density bipolar electrograms from 106 patients (paroxysmal n≈52, non-paroxysmal n≈54) undergoing left atrial mapping with a 24-bipole HD-Grid catheter at standardized sites (RENEWAL AF-ANZCTR ACTRN12619001172190)). Multifractal analysis using the Wavelet Transform Modulus Maxima Method (WTMM) was applied to a burst-energy observable to derive log-normal multifractal parameters c₀ (support dimension), c₁ (spectrum location), and c₂ (fluctuations). Hierarchical mixed-effects models accounted for channels nested within locations within patients. A flecainide sub-study (n=15) provided paired pre-/post-infusion recordings, and 27 spontaneous termination events in 15 patients were analysed using 60-s pre-termination windows. Spatial texture of c₂ was quantified by variogram-derived correlation length and sill.

**Results:** AF electrograms exhibited robust multifractality confirming multifractal fluctuations as an intrinsic property of AF. Non-paroxysmal AF showed significantly reduced fluctuations versus paroxysmal AF (c₂: β=–0.01, p=0.001), indicating a paradoxical loss of fluctuations with disease progression. Flecainide selectively increased fluctuations in paroxysmal AF (Δc₂ = +0.04, p<0.01; Δc₀ = +0.06, p<0.01) but had no significant effect on fluctuations (c2) in non-paroxysmal AF, revealing phenotype-dependent drug response. Immediately prior to spontaneous AF termination, fluctuations increased significantly compared with sustained AF (c₂: 0.198 vs 0.181, p=0.024). Spatial variogram analysis revealed heterogenous patterns in paroxysmal AF, whereas non-paroxysmal AF displayed a homogenised, flattened fluctuations landscape.

**Conclusions:** Atrial fibrillation exhibits robust multifractal dynamics rather than random electrical activity. Reduced fluctuations characterizes non-paroxysmal AF, whereas higher fluctuations is observed in paroxysmal AF, during flecainide modulation, and immediately prior to spontaneous termination. These findings suggest that multifractal fluctuations (c₂) reflects the dynamical state of AF and may serve as a quantitative biomarker of disease progression, pharmacologic responsiveness, and proximity to termination.

**CONDENSED ABSTRACTT:** Atrial fibrillation (AF) exhibits multifractal electrogram fluctuations that vary with disease stage, pharmacologic responsiveness, and proximity to spontaneous termination. In this study, multifractal fluctuations (c₂) was higher in paroxysmal than non-paroxysmal AF, increased selectively with flecainide in paroxysmal AF, and rose immediately before spontaneous termination. These findings identify c₂ as a quantitative marker of AF progression, and imminent reorganization. Clinically, multifractal analysis may enhance intra-procedural assessment of AF phenotype, guide drug selection, and improve recognition of transitions toward sinus rhythm, and connects AF with concepts of criticality and phase transitions.

## Introduction

Atrial fibrillation (AF) is the most common sustained cardiac arrhythmia(2, 3) and a leading cause of stroke, heart failure, and hospitalization worldwide.(3) Despite advances in ablation,(4) important clinical questions remain unresolved, such as methods to reliably distinguish early from advanced AF, relevant to the timing of ablation, the ability to predict treatment response, or identification intra-procedurally of imminent AF termination.

During AF ablation procedures, a familiar observation for clinical electrophysiologists is to see bipolar electrograms momentarily sharpen and organize, then fragment and become fractionated. In this study, we reasoned that these fluctuations, may represent fundamental properties of the AF dynamics that could be used to characterize AF phenotype and progression, response to treatment, and proximity to AF termination. We reasoned that if tools could be developed to systematically capture and quantify these commonly observed shifts in electrogram organization, they could both inform clinical practice and mechanistic understanding of AF.

To date, existing signal analysis approaches used in clinical AF electrogram studies such as dominant frequency,(5) voltage,(1) and entropy(6) analysis and have typically used temporal averaging that smooths over these transient fluctuations in the level of electrogram organization. Here, we hypothesized that these fluctuations, rather than being noise, may represent fundamental properties of the AF dynamics, signatures of self-organization within a complex system, poised near a critical point,(7) attempting to reconfigure toward sinus rhythm. The brief and non-stationary nature of these fluctuations makes them difficult to quantify using time-averaged electrogram metrics. Multifractal analysis is designed to capture short-lived, burst-like changes in signal organization across temporal scales that has previously been used in surface ECG analyses to discriminate disease states,(8) but has previously only received limited examination in AF.(9) We hypothesized that the same approach applied to intracardiac AF electrograms could provide clinically actionable markers of phenotype, drug responsiveness, and imminent termination during procedures. The demonstration that AF is multifractal in character would have significant mechanistic implications, as a marker of AF dynamics indicating proximity to a critical phase transition.(7)

## Methods

### Study Population and Data Acquisition

We analysed high-density left atrial electrograms from 106 patients with atrial fibrillation (AF) undergoing clinically indicated mapping procedures (52 paroxysmal, 54 non-paroxysmal) according to a pre-specified protoocol.(10) Patients were recruited under Southern Adelaide Local Health Network Human Research Ethics Committee approval (282.19), and all provided written informed consent. Inclusion criteria were age ≥18 years and documented AF at the time of mapping. Exclusion criteria included prior left atrial surgery or inability to maintain catheter stability during mapping.

Electrograms were recorded using a 24-channel HD-Grid catheter (Abbott, USA) connected to the EnSite mapping system (EnSite Precision or EnSite X). Raw electrograms were sampled at 2,000 Hz with 50Hz notch and bandpass filters set by the EnSite System (30Hx-500Hz). The catheter was positioned sequentially at ten standardized left atrial sites: anterior wall, septum, high posterior, low posterior, left atrial appendage, lateral wall, and the ostia of the left and right superior and inferior pulmonary veins. Contact was confirmed by electroanatomic mapping and electrogram quality. At each site, a ≥60-second segment was acquired using the “segment recordings” feature on the mapping system. In a subset of 15 patients (6 paroxysmal, 9 non-paroxysmal), paired recordings were obtained before and after intravenous flecainide infusion during posterior wall mapping. Baseline recordings were acquired immediately before infusion, and repeat recordings were obtained after approximately 400 seconds of continuous infusion.

All recordings were screened for catheter stability and signal quality by an independent electrophysiologist. A custom software platform (WaveWrangler, Flinders University, Australia) was used to filter and extract bipolar electrogram data from each location segment/recording. De-identified binary files (.npy) with accompanying coding metadata file were exported for data analysis. All analysis code was written in python programming language using open source python extension libraries, and executed using Jupyter notebook Python 3 (**Python 3.12.9** (packaged by conda-forge; main, Feb 14 2025, 08:00:06).

### Signal Processing

Electrogram recordings were decimating to 500 Hz and cubic-spline differentiation and squaring to generate the burst-energy signal *E*(*t*) was computed(9). Dynamic fluctuations were quantified using a continuous wavelet transform with a third-order Gaussian derivative wavelet, analysing scales corresponding to physiologic periodicities from 0.6–10 s. Local maxima of the wavelet modulus, representing energy bursts across scales were identified as the basis for constructing scale-dependent partition functions. Scaling exponents τ(*q*) were obtained from log–log regressions and fitted to a log-normal multifractal model to derive parameters including *c*_2_, a coefficient reflecting the intensity of multifractal electrical fluctuations.

To assess the spatial organization of *c*_2_ across high-density maps, we constructed empirical semi-variograms using electrode midpoints and Euclidean distances between all bipolar pairs, with lags binned at 4 mm consistent with HD-Grid geometry. Analysis was constrained to the physical measurement footprint (0–17 mm) to avoid model-based extrapolation, and an exponential variogram model was fitted to estimate two clinically interpretable spatial parameters: the sill, representing the total magnitude of local heterogeneity, and the range, representing the correlation length over which tissue-level organization is preserved. Together, these temporal and spatial multifractal metrics provide a quantitative framework for characterizing the complexity and texture of atrial electrical activity in AF.

## Statistical Analysis

### Data Structure and Hierarchical Modelling

Continuous variables were expressed as mean ± standard deviation (SD) or median where appropriate. Categorical variables were compared using the Chi-square or Fisher’s exact test. To account for the hierarchical structure of the high-density mapping data (where multiple bipolar electrogram channels are nested within specific anatomical locations, which are in turn nested within individual patients), linear mixed-effects models were employed. These models included random intercepts for both patient and location to adjust for intra-cluster correlation.

### Phenotype and Chamber Comparisons

The primary analysis compared multifractal parameters (c_0_, c_1_, c_2_) between paroxysmal and non-paroxysmal AF phenotypes. Fixed effects in the mixed-effects models included AF phenotype and atrial chamber (left vs. right atrium). Interaction terms were tested to determine if phenotype-specific differences were consistent across chambers. Variance partitioning was performed to estimate the proportion of total multifractal variability attributable to patient-level differences, local anatomical heterogeneity (location-level), and residual channel-level variability.

### Intervention and Termination Analysis

For the flecainide sub-study, paired pre- and post-infusion recordings were compared within patients using paired t-tests or Wilcoxon signed-rank tests, depending on data distribution. Interaction analysis was utilized to evaluate whether the pharmacologic response of multifractal signatures differed significantly by AF phenotype. For spontaneous termination events, mixed-effects models compared the 60-second pre-termination window against segments of ongoing AF, adjusting for phenotype and clustering.

### Spatial Statistics

The spatial organization of fluctuations (c_2_) was quantified using empirical semi-variograms. Lags were binned at 4 mm intervals to align with the HD-Grid catheter geometry, with analysis constrained to the 17 mm physical measurement footprint to avoid extrapolation. An exponential variogram model was fitted to estimate the sill (total magnitude of local heterogeneity) and the range (correlation length).

### Software and Significance

All statistical analyses were performed using Python 3.12 with open-source extension libraries. A p-value of < 0.05 was considered statistically significant for all tests.

## Results

### Baseline Characteristics of the Clinical Cohort

Baseline characteristics of the cohort are summarised in Table 1. A total of 106 patients (mean age 60.7 ± 10.5 years) undergoing catheter ablation for atrial fibrillation (AF) were enrolled in the study. The cohort was stratified by clinical phenotype into paroxysmal AF (n=52) and non-paroxysmal AF (n=54) groups. The non-paroxysmal cohort demonstrated clinical markers of more advanced atrial and ventricular remodelling compared to the paroxysmal group. Patients with non-paroxysmal AF had significantly larger left atrial diameters (70.4 ± 19.5 mm vs 59.8 ± 17.5 mm, p=0.035) and a higher prevalence of heart failure (54% vs 22%, p < 0.001). Left ventricular ejection fraction was significantly lower in the non-paroxysmal group (55.3 ± 11.9% vs 60.5 ± 9.1%, p < 0.001). Primary demographics like age (p=0.44), male sex distribution (p=0.89), and BMI (p=0.06) were balanced between the groups. Non-paroxysmal group had a higher utilization of Class Ic antiarrhythmic medications (42% vs 14%) at the time of the procedure. These findings establish that the non-paroxysmal cohort represents a more entrenched disease state characterized by structural and functional substrate progression, providing a clinically relevant basis for comparative analysis.

**Table 1.** Baseline Demographics, Comorbidities, and Echocardiographic Parameters Categorized by AF Phenotype.

| Variable | Paroxysmal AF (n=52 ) | Non-Paroxysmal<br>AF (n= 54) | p-value |
| --- | --- | --- | --- |
| Age | 60.53 ± 11.4 years | 60.83 ± 9.7 years | 0.44 |
| Male sex | 34 male (65%) | 46 male (85%) | 0.89 |
| BMI (kg/m<sup>2</sup>) | 29.2 ± 4.4 | 30.7 ± 5.5 | 0.06 |
| Hypertension | 26 (51%) | 19 (36%) | 0.18 |
| Diabetes mellitus | 5 (10%) | 6 (11%) | 0.9 |
| Heart failure | 11 (22%) | 30 (54%) | <0.001 |
| Left ventricular ejection fraction<br>(%) | 60.5% ± 9.1% | 55.3% ± 11.9% | <0.001 |
| Left atrial diameter (mm) | 59.8mm ± 17.5mm | 70.4mm ± 19.5mm | 0.035 |

### Atrial Fibrillation Bipolar Electrograms Are Multifractal

Multifractal analysis of 27,869 bipolar electrograms from 1,990 sites across 106 patient left atria revealed robust scale-invariant complexity in AF signals that deviated substantially from monofractal behaviour (Figure 4). The distribution of Hölder exponents demonstrated a consistent hierarchical ordering, with C₀ (mean 1.130 ±0.119) > C₁ (mean 0.729±0.164) > C₂ (mean 0.203±0.066) observed in 99.1% of recordings (Figure 4 Panel A). This systematic ordering indicates that AF electrograms contain a spectrum of singularity strengths rather than uniform temporal scaling, with C₂ representing the strongest, most irregular fluctuations in the signal.

**Figure 1.**
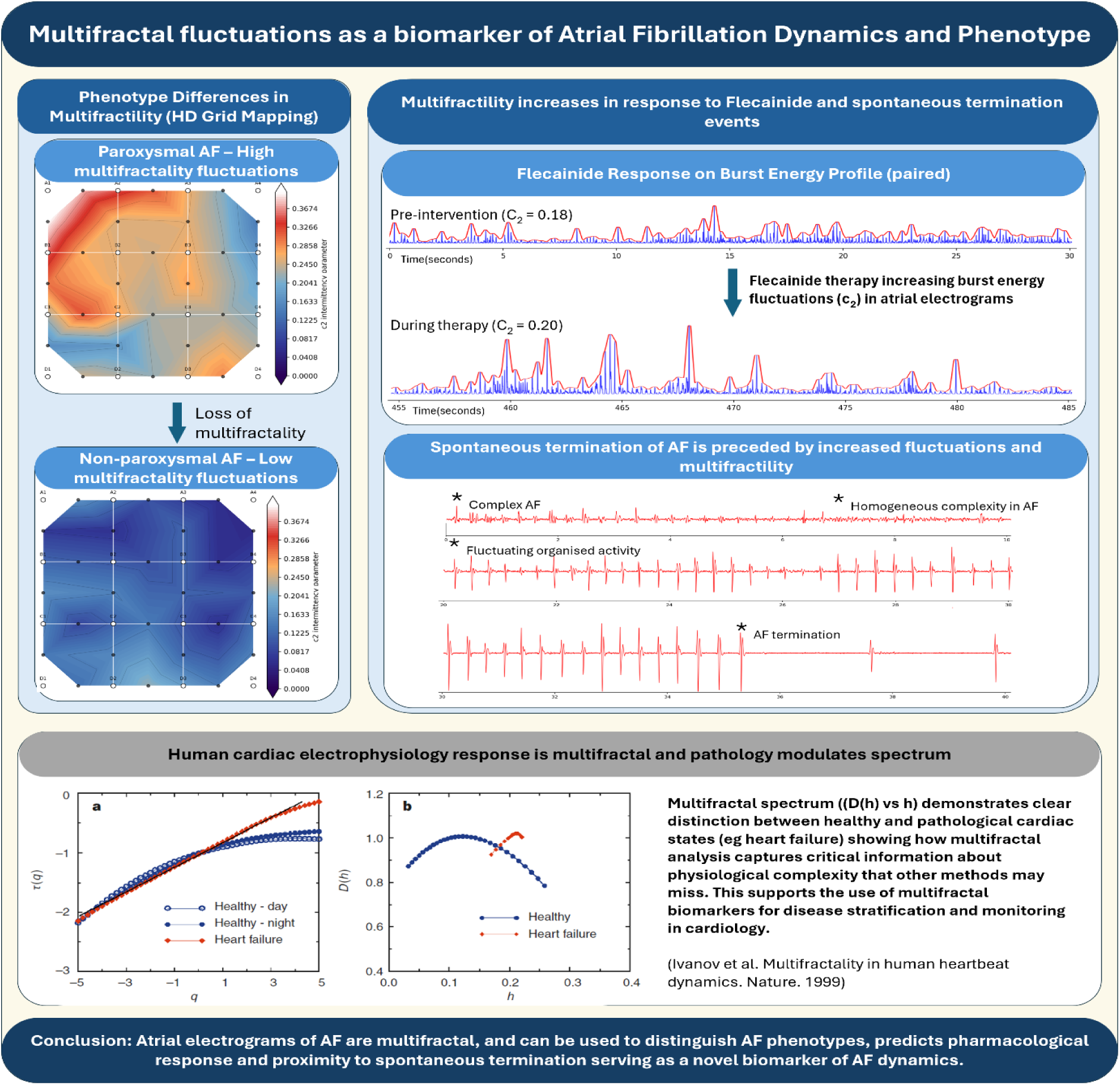
Graphical Abstract: Left: High-density mapping demonstrates greater multifractal fluctuations (c₂) in paroxysmal AF, characterized by spatial heterogeneity and burst-like electrogram fluctuations, compared with non-paroxysmal AF, which exhibits reduced fluctuations and a spatially homogenized landscape. Top right: Intravenous flecainide increases burst-energy fluctuations and multifractal fluctuations (c₂) in paroxysmal AF, indicating preserved dynamical range, whereas non-paroxysmal AF shows minimal response. Bottom right: Spontaneous AF termination is preceded by increased electrogram fluctuations and elevated fluctuations, suggesting proximity to reorganization. Bottom: Adapted from Ivanov et al (Nature 1999) demonstrating multifractal spectra scaling properties distinguish physiological from pathological cardiac dynamic in heart failure

**Figure 2.**
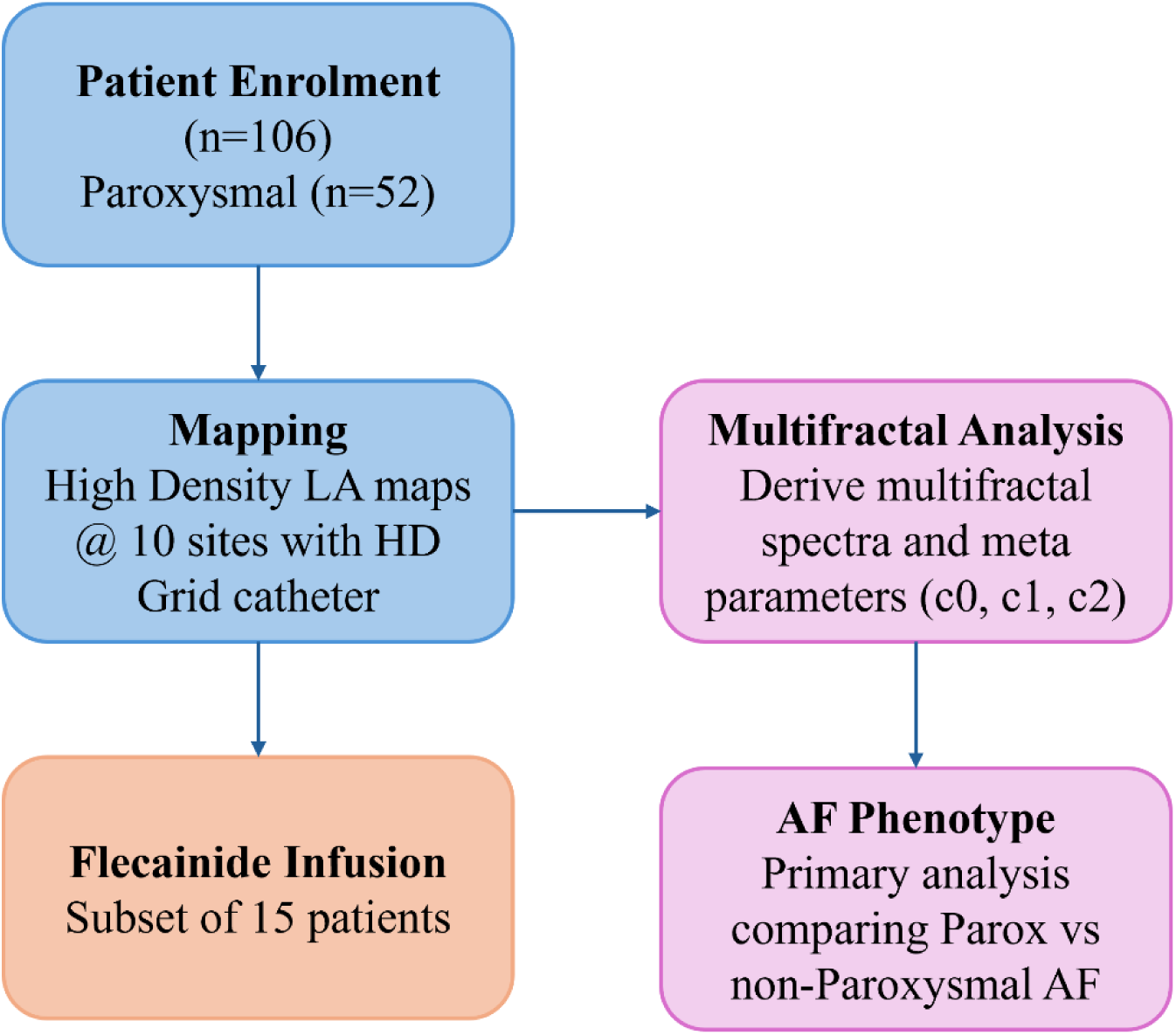
Study workflow for multifractal analysis of atrial fibrillation (AF). Patients with AF (n = 106; 52 paroxysmal, 54 non-paroxysmal) underwent high-density left atrial mapping using a 24-channel HD-Grid catheter at ten standardized sites. Electrograms were processed using the Wavelet Transform Modulus Maxima (WTMM) method to derive multifractal parameters (c₀, c₁, c₂). Primary analyses compared dynamic electrical propagation (c₂) between AF phenotypes at chamber, regional, and local scales. In a subset of 15 patients, paired recordings before and after intravenous flecainide infusion were analysed to assess pharmacologic modulation of multifractal properties.

**Figure 3.**
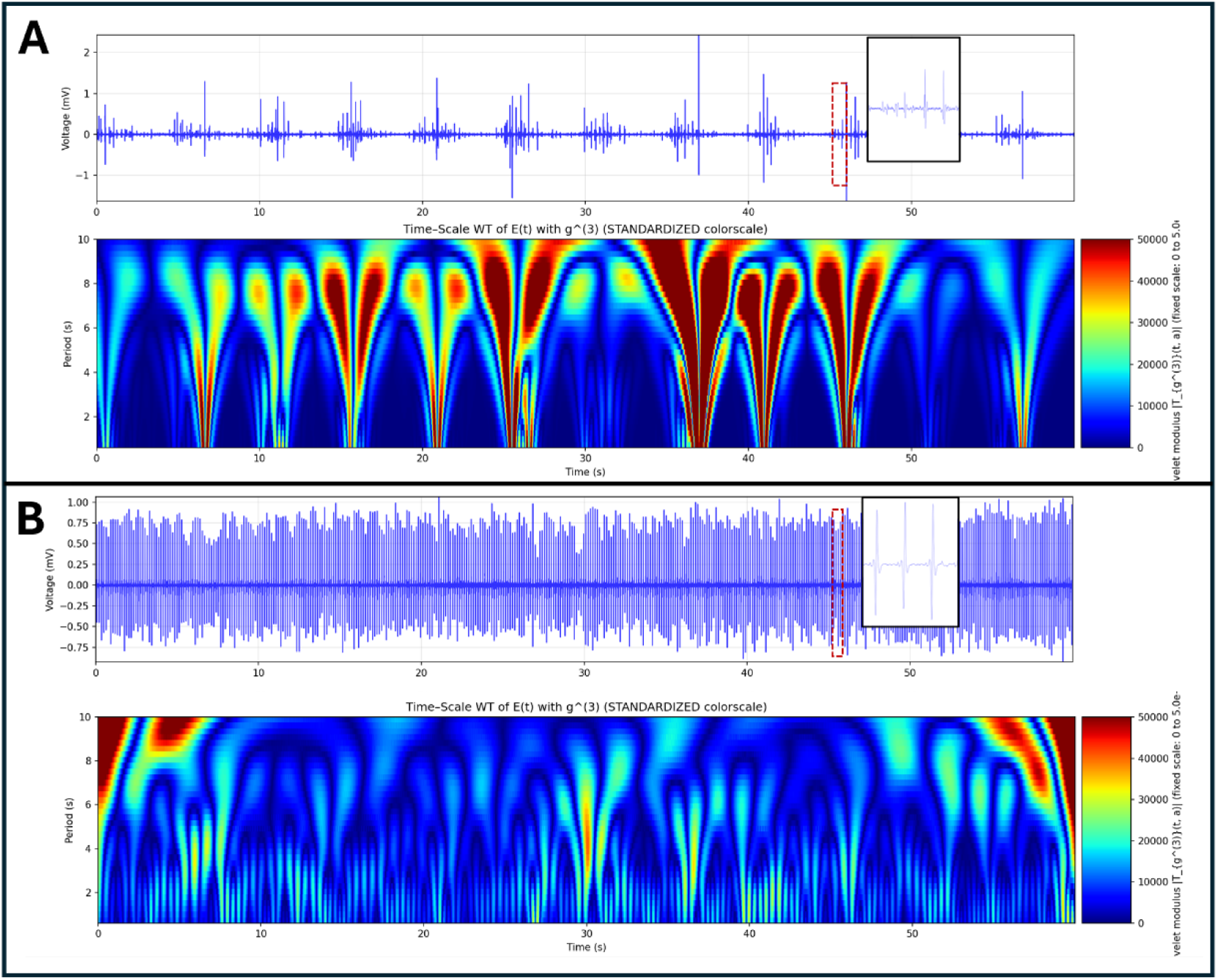
Electrogram and corresponding multifractal spectral heat map of atrial fibrillation electrogram dynamics in two representative patients; A – recording showing a period of high intermittent changes in electrogram, and B, during a period of high organisation. (A): Upper frame showing raw bipolar electrogram signal over 60 seconds showing intermittent changes in electrograms throughout. Lower frame showing the corresponding wavelet transform scalogram of E(t) heatmap with high fractal energy during intermittent changes in electrogram across the spectra across all wavelet periods indicating multifractal energy dynamics. (B): Upper frame showing raw bipolar electrogram signal over 60 seconds showing temporally organised electrograms throughout. Lower frame showing corresponding wavelet transform scalogram of E(t) heatmap showing lower fractal energy throughout, with changes in energy across lower wavelet periods (bands of light blue and dark blue) at lower wavelet periods.

**Figure 4.**
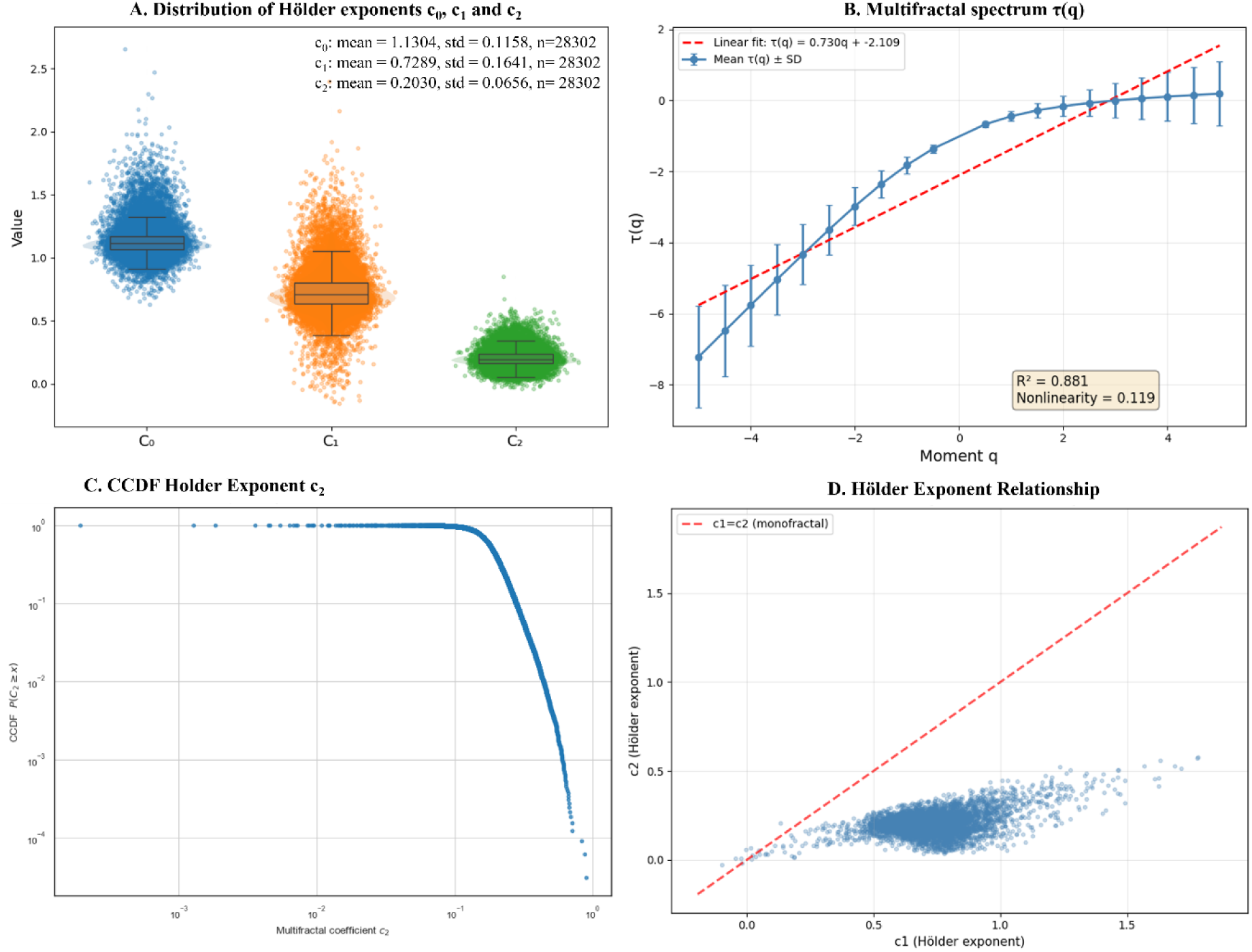
Mulitfractal analysis of n=27,869 bipolar electrograms from 1,990 sites in left atria during sustained Atrial Fibrillation from 106 patients. A. Distribution of Hölder exponents C₀, C₁, and C₂. Boxplots are overlaid with jittered scatter points. Summary statistics: C₀ ( mean=1.1304±0.1158), C₁ (mean=0.7289±0.1641), C₂ (mean=0.2030±0.0656). The ordering C₀ > C₁ > C₂ (99.19% of recordings) indicates clear multifractal scaling in AF electrograms. Note C₂ representing the strongest singularities. B. Scaling function τ(q) versus moment q. The nonlinear relationship between τ(q) and q, indicating strong multifractal scaling. The solid line represents the fitted curve, while the dashed line shows the linear monofractal reference (R² = 0.88). C. Complementary Cumulative Distribution Function (CCDF) of the fluctuations parameter c2 on a log-log scale. The distribution exhibits a heavy tail characteristic of complex dynamical systems D. Relationship between Hölder exponents c1 and c2. Scatter plot illustrating the correlation between the Hölder exponents c1 and c2 across a random sample of 5,000 recordings from 27,869 total electrograms. The red dashed diagonal line represents the monofractal reference (c1 = c2), where multifractal signals typically exhibit c1 > c2. The distribution below the diagonal confirms multifractal scaling in AF signals, with c1 and c2 showing distinct values indicative of complex temporal dynamics.

**Figure 5.**
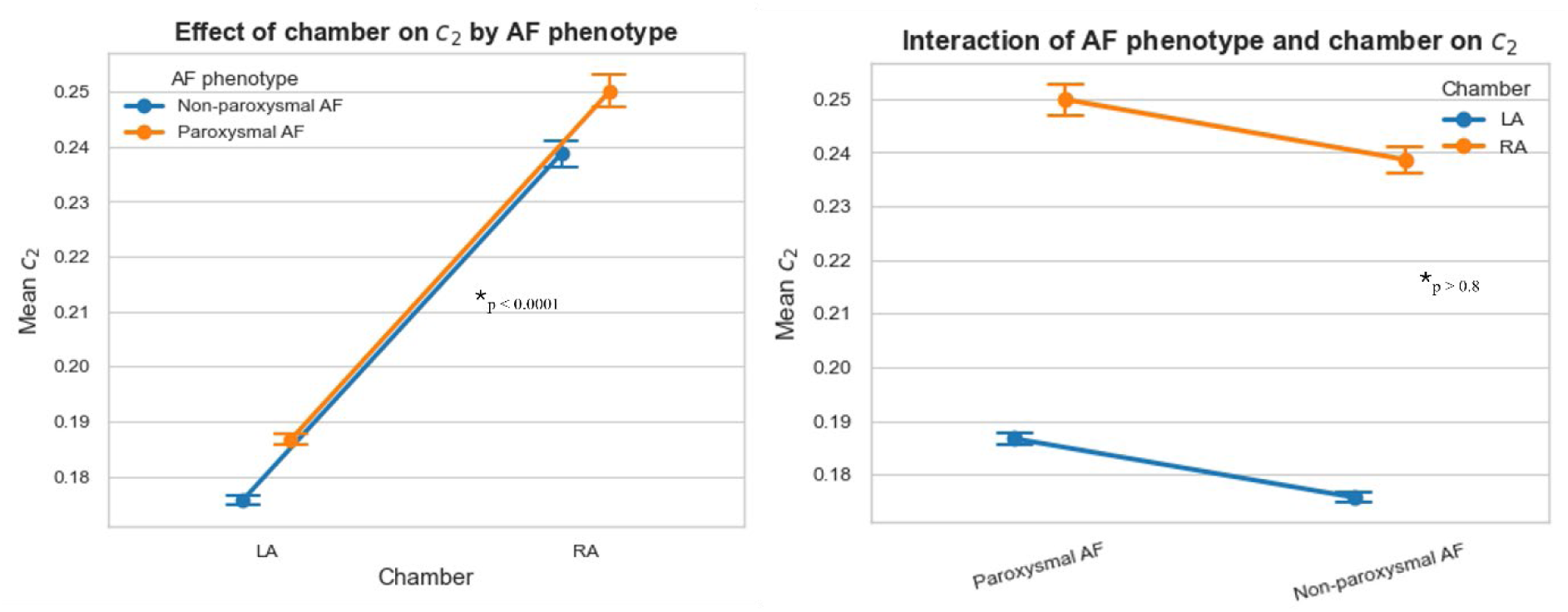
There were significant differences in the c2 parameter between RA and LA (p<0.001), however these differences were consistent across AF phenotype with no interaction between chamber and phenotype (p>0.4)

The scaling function τ(q) exhibited marked nonlinearity with respect to moment order q, diverging substantially from the linear monofractal reference (R² = 0.88, Figure 4 B). This nonlinear relationship confirms that different temporal scales in AF electrograms display distinct scaling behaviours, inconsistent with simple fractal or stationary models, and most consistent with a multifractal spectral model.

The distribution of multifractal spectral width (C₀ - C₂) showed a mean width of 0.928, substantially exceeding the monofractal reference of zero, demonstrating that AF electrograms require multiple indices to capture the full spatiotemporal complexity.

The relationship between C₁ and C₂ further confirmed multifractal scaling, with the majority of recordings distributed below the monofractal diagonal reference line (C₁ = C₂) (Figure 4 D). The spatial separation between C₁ and C₂ values reflects the coexistence of multiple temporal patterns within individual electrograms, consistent with intermittent switching between organized and disorganized activation sequences. This separation persisted across the entire dataset, indicating that multifractal structure is an intrinsic and reproducible property of AF electrograms across anatomical locations and AF phenotype. Furthermore, to characterize the statistical nature of these fluctuations, we analysed the distribution of the fluctuations parameter (C2). The data exhibited a heavy-tailed distribution that was best described by a Log-Normal model (p < 10^-5^) (Figure 4 C).

### RA vs LA differences

Across all recordings, mixed-effects models with random intercepts for case and case–location showed a robust chamber effect on all multifractal coefficients. Using paroxysmal AF as the reference and left atrium (LA) as the reference chamber, mean c₀ in LA paroxysmal AF was LA c0 = 1.09 vs RA c0 = 1.23; p < 0.001) and a small but significant reduction in non-paroxysmal AF (p = 0.007). Similarly, c₁ was higher in RA than LA (p < 0.001), with no clear effect of AF phenotype (p = 0.61). For c₂, LA paroxysmal AF was significantly higher in c2 than RA (LA c2= 2.05 vs RA c2= 0.19, p < 0.001), indicating consistently greater fluctuations in the right atrium. Non-paroxysmal AF showed lower c₂ than paroxysmal AF for C2 (p = 0.005), however the interaction between RA and LA with phenotype were non-significant for all three Holder coefficients (p > 0.4). These results suggest that multifractal LA–RA differences are stable across AF phenotypes rather than phenotype-specific.

### Multifractal Spectra Significantly Differ Between Paroxysmal and Non-paroxysmal AF in the LA

As the differences between LA and RA were equivalent between AF phenotype, initial assessment excluded RA locations from analysis. After WTMM on all left atrial electrograms, both paroxysmal and non-paroxysmal AF demonstrated clear multifractal scaling behaviour with non-linear spectral scaling τ(q) curve [Figure 6 A]. Linear mixed-effects models that accounted for the hierarchical structure of electrograms nested within locations within patients, non-paroxysmal AF was associated with statistically significant reductions in the Hölder exponents c₀ (β = –0.019, 95% CI –0.031 to –0.008, p = 0.001) and the fluctuations parameter c₂ (β = –0.011, 95% CI –0.018 to –0.004, p = 0.001). In contrast, c₁ showed no significant difference between AF phenotypes (β = –0.007, 95% CI –0.035 to 0.020, p = 0.590)[Figure 6 C].

**Figure 6.**
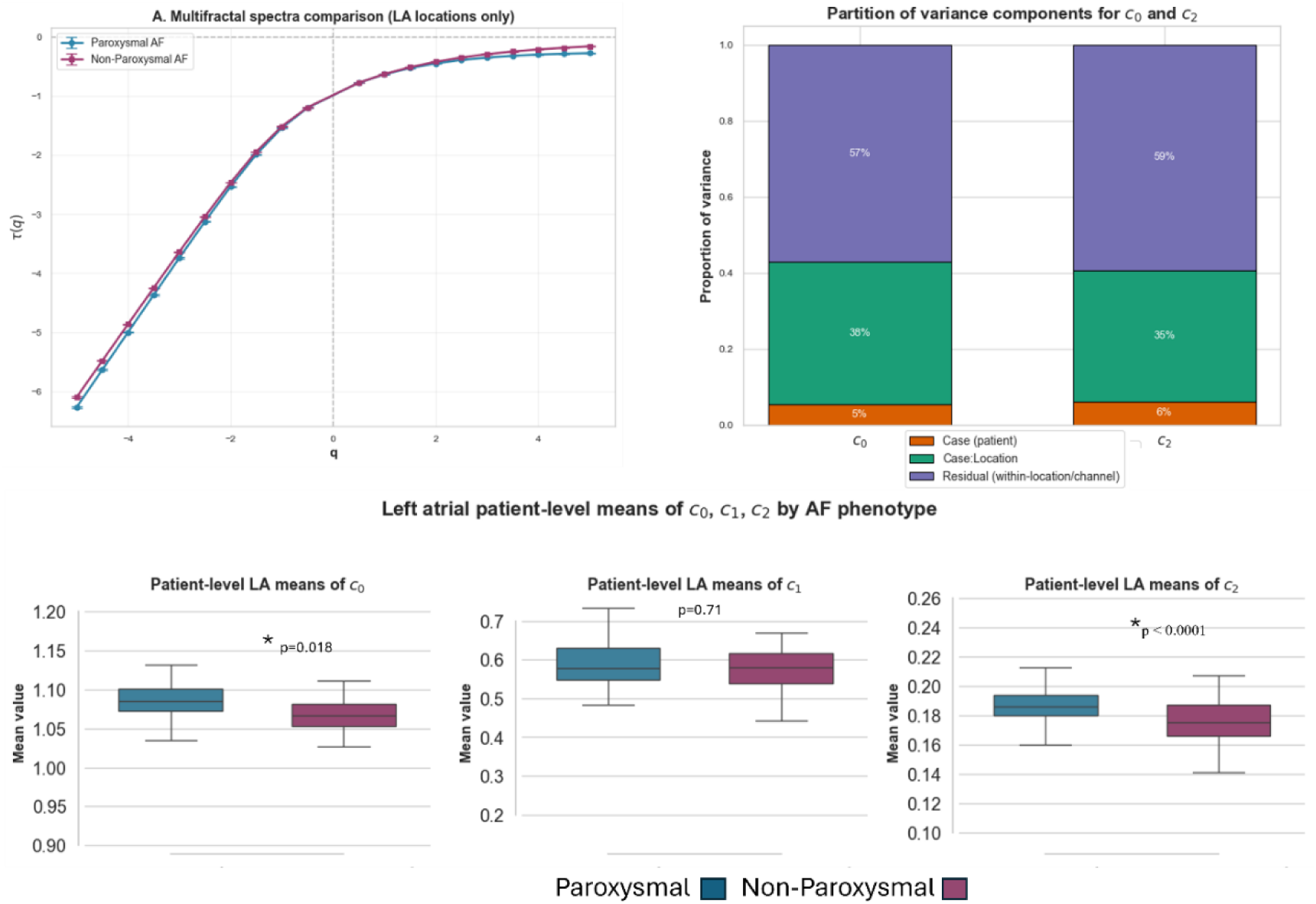
**A:** Multifractal spectra comparison between paroxysmal and non-paroxysmal AF, with both phenotypes demonstrating multifractal scaling, with increased spectral width in paroxysmal AF compared to non-paroxysmal patients. **B: P**roportion of total variance in Holder exponent C0 and C1 attributable to differences between patients (Case), between recording locations within patients (patient: Location), and within-location/channel-level residual variability (Residual), based on the mixed-effects models. For c0, only a small fraction of variability arises at the patient level (∼5%), with much larger contributions from location-level (∼38%) and residual (∼57%) variance. For the fluctuations parameter *c*_2_, patient-level variance is negligible, with variability dominated by location-level (∼35%) and residual (∼59%) components. These variance partitions indicate that multifractal dynamics are governed primarily by local spatial heterogeneity within the atria, rather than by global differences between patients such as AF phenotype or other degrees of disease state. **C:** Boxplots of left atrial multifractal Holder exponent showing significant differences in c0= and c2, with no difference in C_1_.

**Figure 7.**
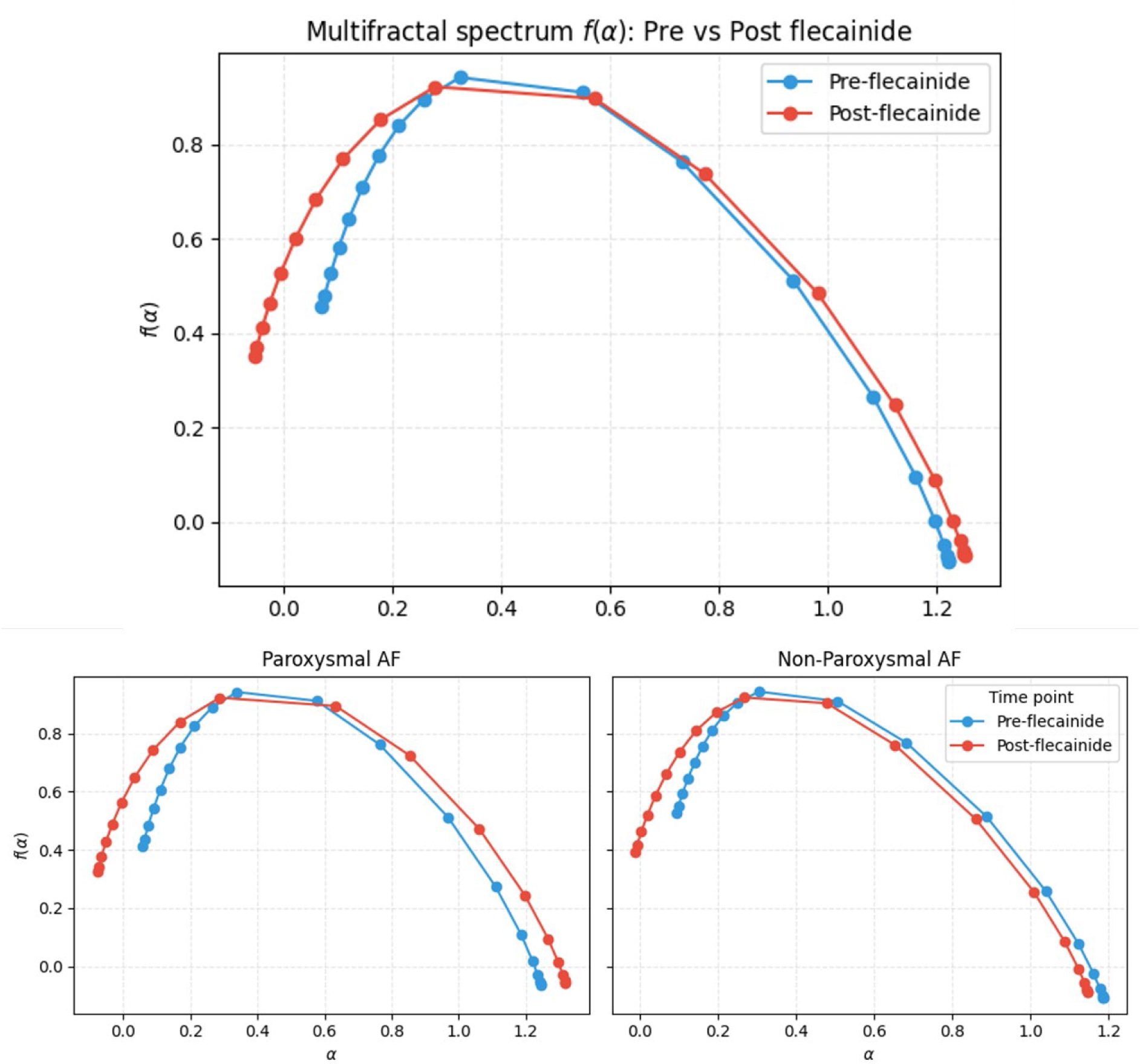
Multifractal spectra of atrial electrograms before (blue) and after (red) flecainide infusion. Charts show multifractal spectrum (f(α)) as a function of the Hölder exponent (α). Spectra were derived from (τ(q)) estimates over a wide range of moments and averaged across all 60-s segments and channels within each group. **Top:** Overall effect of Flecainide on multifractal spectra showing increase spectral width with after flecainide infusion. **Lower:** (left) paroxysmal AF and (right) non-paroxysmal AF spectral changes due to flecainide infusion.

**Figure 8.**
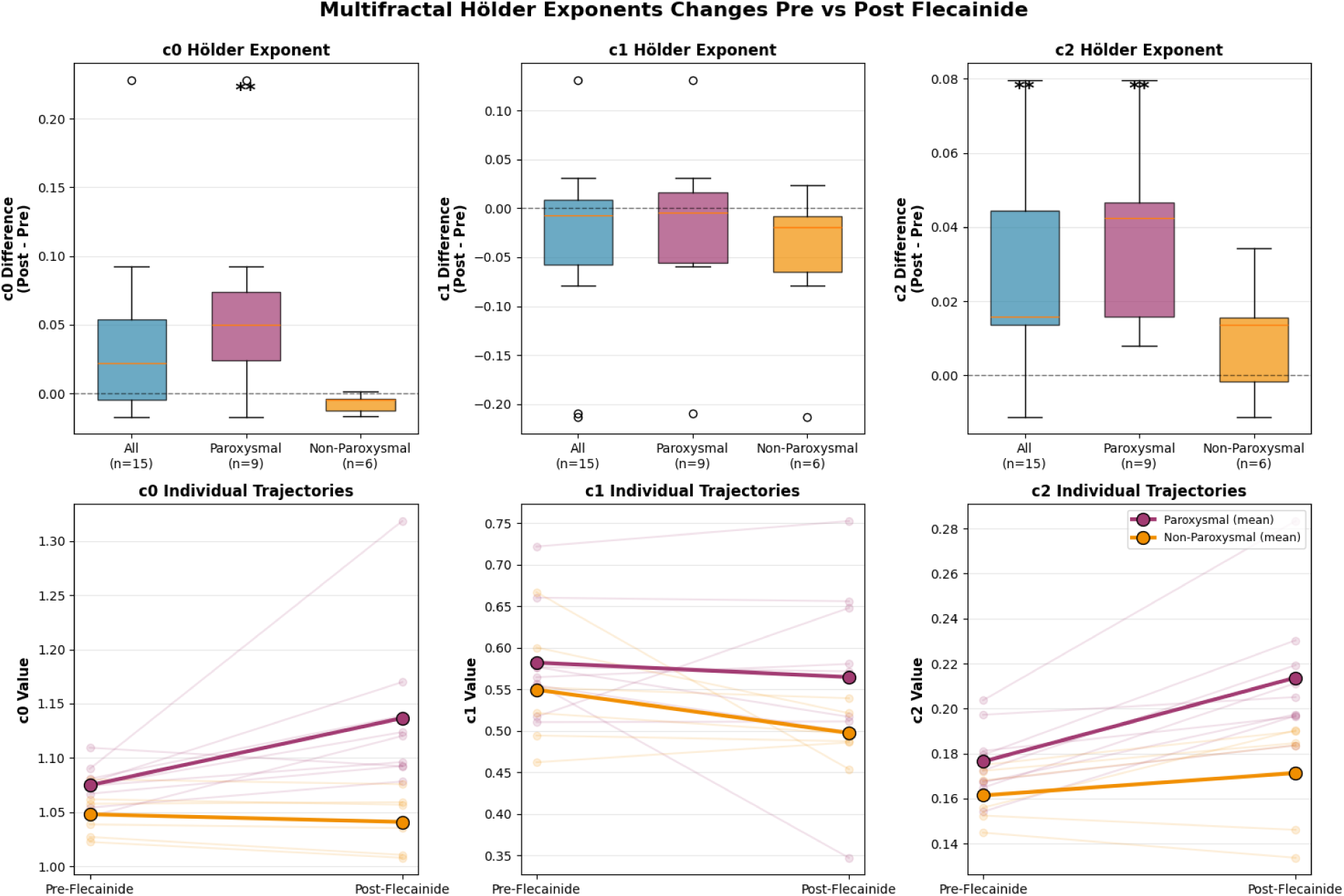
Changes in multifractal Hölder exponents (c0, c1, c2) before and after flecainide administration in patients with atrial fibrillation. **TOP:** Flecainide significantly increased the long-range temporal organisation (c0) of AF in paroxysmal patients (p<0.001) with no limited affect (p=0.0625) in non-paroxysmal patients. Flecainide and no effect on the fractal dimension (c1) Holder exponent for either paroxysmal (p=0.6523 or non-paroxysmal AF (p=0.1562). Flecainide significantly increased the fluctuations of electrograms in paroxysmal patients (p < 0.01), however did not affect the fluctuations in non-paroxysmal patients (p = 0.1563). **Lower:** displays individual patient trajectories with group mean lines. Significant changes (Wilcoxon signed-rank test, p < 0.05) are indicated by asterisks. These findings suggest flecainide modulates multifractal properties differentially based on disease phenotype. AF.

Mixed effect variance partitioning demonstrated that multifractal variability was dominated by location-level heterogeneity, with the Patient: Location random-effect variance exceeding the between-patient variance for all parameters (c₀: 0.003 vs 0.001; c₁: 0.011 vs 0.004; c₂: 0.001 vs 0.000). This indicates that spatial variability in local electrogram dynamics across the grid catheter is the dominant contributor to the differences observed within patients [Figure 6 B].

### Flecainide Modulates Multifractal Spectra in Paroxysmal AF

In a sub-group of 15 patients (9 paroxysmal, 6 non-paroxysmal AF), we intervened in the AF dynamics with intravenous flecainide while the HD grid catheter was placed on the posterior left atrium. Sixty seconds of grid bipolar electrograms were exported prior to infusion, and another 60 seconds were recorded immediately after infusion (mean time between recordings Δt =506.38 seconds, 95% CI 366.10 to 646.65 seconds). All electrograms were analysed with the same parameters for WTMM methods as described above. Overall analysis revealed a significant increase in multifractal spectra [Figure 9, Top] with holder exponent c_2_ (fluctuations) showing greatest increase during flecainide infusion (Δ c_2_ = +0.026, p <0.001), while changes in global scaling (Δc_0_ = +0.034, p = 0.064) and fractal dimension (Δc1 = −0.031, p = 0.188) did not reach statistical significance (Figure 10).

**Figure 9.**
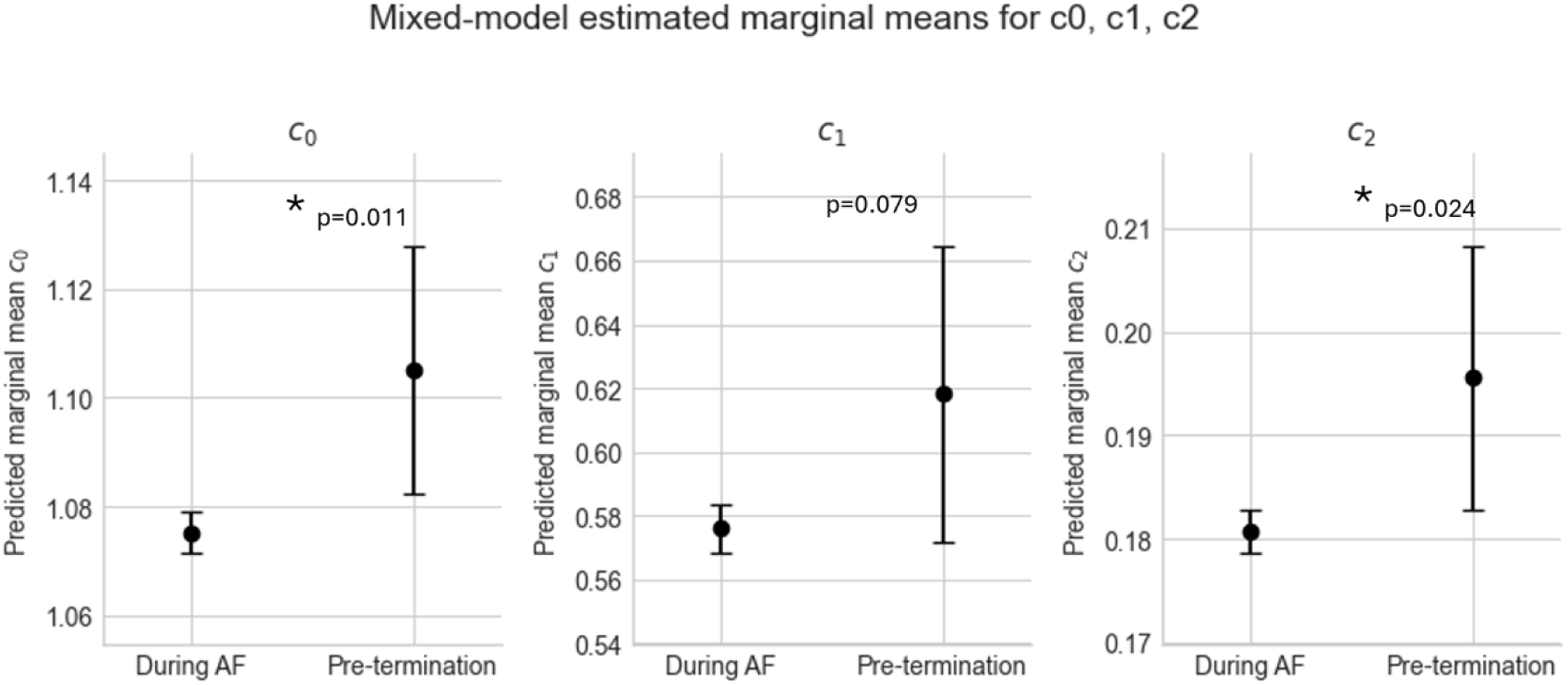
Predicted marginal mean of Holer Exponents c0, c1, c2) during ongoing atrial fibrillation and in the 60 s pre termination window, estimated from a linear mixed effects model (c2 ∼ AF_Status_bin + AF_Type_bin + (1|Case) + (1|Case:Location)). Points show fixed effect marginal means with 95% confidence intervals, demonstrating a small but statistically significant increase in c₂ immediately prior to AF termination compared with ongoing AF.

**Figure 10.**
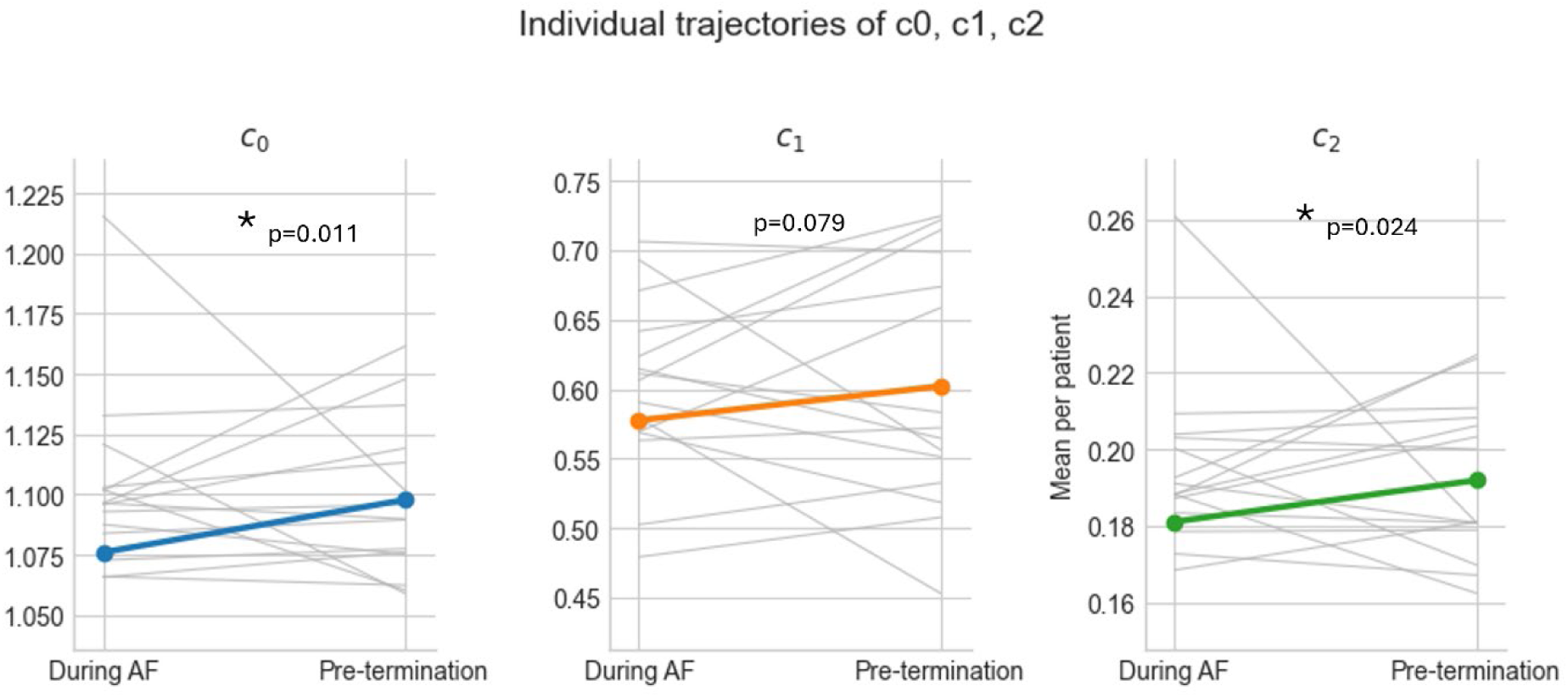
Individual patient trajectories of Holder exponents c₀, c₁, and c₂ from ongoing atrial fibrillation recordings (“During AF”) to the 60 s pre-termination window (“Pre-termination”). Thin grey lines show each patient’s mean value at both time points, while the coloured lines (blue for c₀, orange for c₁, green for c₂) indicate the corresponding population mean changes, highlighting increase in Holder exponents c0 can c2 prior to AF termination.,

**Figure 11.**
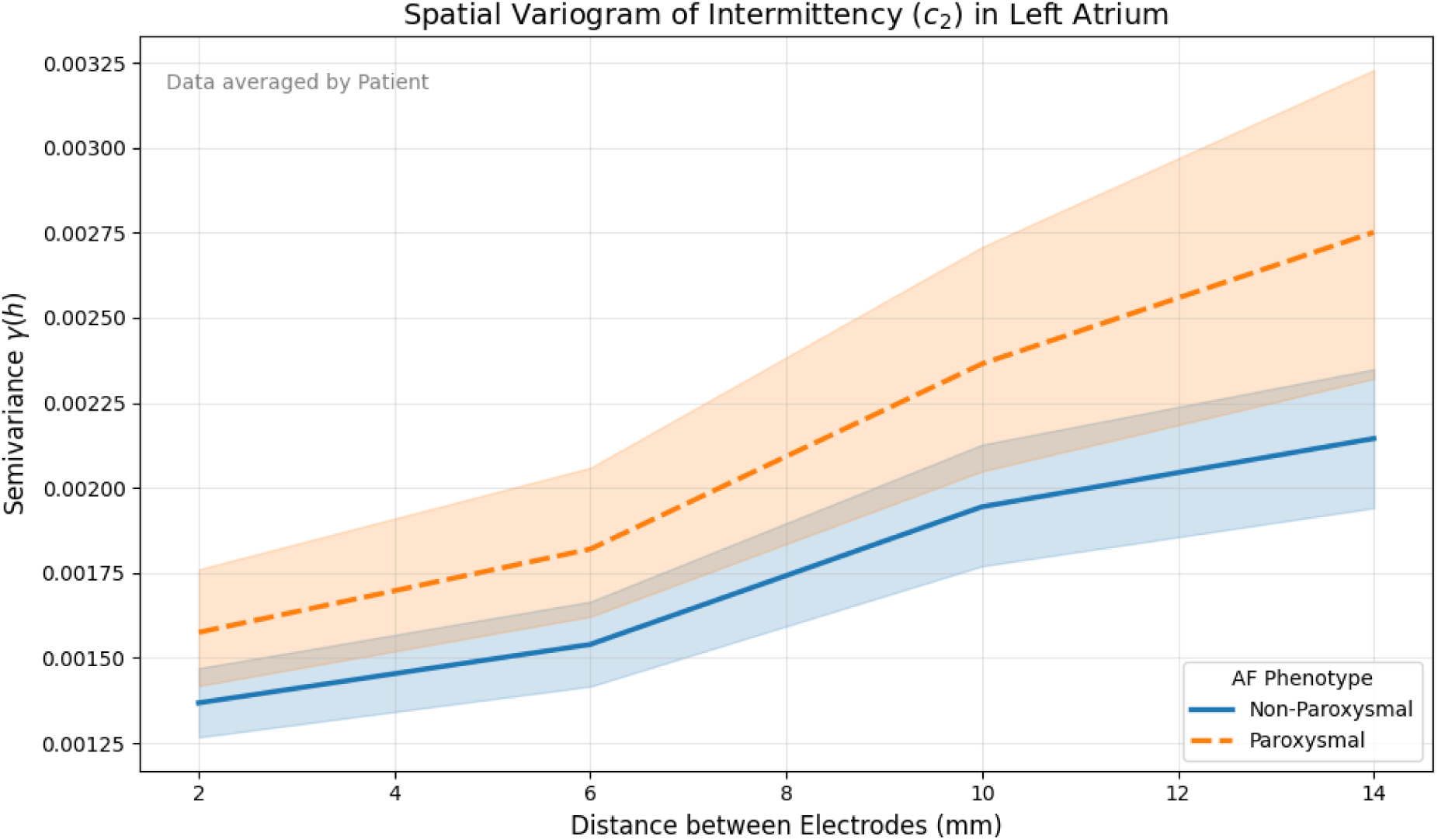
Spatial Variogram of Fluctuations (c_2_) in the Left Atrium. The plot displays the patient-averaged semi-variance (γ(h)) of the fluctuations parameter c2 as a function of distance between electrode bipoles (n=106 patients). **Orange (Dashed):** Paroxysmal AF (n=52) demonstrates higher semi-variance, indicating a spatially heterogeneous (“patchy”) substrate where dynamical properties vary significantly over short distances. **Blue (Solid):** Non-Paroxysmal AF (n=54) demonstrates a flattened curve with lower semi-variance, indicating a homogenized substrate with uniformly suppressed fluctuations. **Shaded regions** represent 95% confidence intervals. The rising slope without a plateau indicates that the correlation length of multifractal dynamics exceeds the catheter dimensions (>17 mm) in both phenotypes.

On subgroup analysis by AF phenotype (paroxysmal n=9, non-paroxysmal n=6), the spectral changes observed across all patients was revealed to be driven by the paroxysmal cohort [Figure 9, Lower]. Paroxysmal AF patients demonstrated significant increases in both global scaling (Δc_0_ = +0.062, p = 0.0078) and multifractal width (Δ c_2_: +0.037, p = 0.0039) following flecainide infusion [Figure 10]. In contrast, non-paroxysmal AF patients showed no significant changes in any Hölder exponent (Δc_0_: p = 0.063, Δc_1_: p = 0.156, Δc_2_: p = 0.156), suggesting in-elastic disease state. Fractal dimension (c_1_) remained unchanged in both phenotypes (paroxysmal: p = 0.652; non-paroxysmal: p = 0.156), indicating that flecainide modulates the dynamic range of the electrogram characteristics (more intermittent bursts), while still remaining multifractal in nature.

### Multifractal Signatures of Spontaneous AF Termination Reveal Phenotype-Dependent Reorganization Patterns

During mapping, there spontaneous reversion to sinus rhythm occurred in 27 locations in 15 patients. Terminations events were seen in both AF phenotypes (paroxysmal: 3.89%, n= 19/488 locations; non-paroxysmal: 1.48% terminations, n = 8 /542 locations). For each termination event, 60 seconds of electrograms were exported prior to the event, and were manually assessed to ensure exports did not include sinus rhythm with exports ending <20ms after the last AF electrogram.

Mixed-effects models adjusted for AF phenotype and patient- and location-level clustering was used with results showing higher values of all three Holder exponents in the 60 seconds prior to spontaneous termination compared with ongoing AF segments: (C_0_: 1.109 vs 1.075, p = 0.011), C_1_: 0.620 vs 0.576, p = 0.08, C_2_: 0.198 vs 0.181 respectively, p = 0.024).

### Spatial Organisation of Fluctuations Dynamics

To determine whether the reduction in fluctuations (c_2_) observed in non-paroxysmal AF was a global phenomenon or the result of spatial fragmentation at the catheter location, we performed a variogram analysis. Given the significant inter-chamber differences observed in the primary analysis, this was restricted to the Left Atrium. To ensure statistical independence, variogram parameters were calculated for each map and aggregated to a single mean value per patient (N=106).

### Absence of Micro-Scale Fragmentation

In both phenotypes, the semi-variance of c_2_ (γ(h)) exhibited a continuous rise throughout the observable window of the catheter (0–17 mm) without reaching a plateau. The estimated spatial correlation length for c_2_ (Range) was similar between phenotype (Paroxysmal: 53.8 ± 14.7 mm; Non-Paroxysmal: 52.9 ± 11.6mm) and exceeded the physical footprint of the mapping electrode (12mmx12mm). This indicates fluctuations dynamics are largely un-fragmented at the local (millimeter) scale and may be governed by larger patterns beyond the mapping catheter footprint.

### Loss of Spatial Texture in Non-Paroxysmal AF

Despite similar correlation lengths for fluctuations (c_2_) with AF phenotype, the magnitude of spatial heterogeneity (Sill) differed between groups (Figure 13). Paroxysmal AF exhibited a “rougher” dynamical landscape, characterized by a higher mean semi-variance (0.0049 ± 0.0042) compared to non-paroxysmal AF (0.0036 ± 0.0018).

Notably, the Paroxysmal cohort displayed markedly higher inter-patient variability in spatial texture (σ= 0.0042) compared to the tighter clustering observed in the non-paroxysmal cohort (σ= 0.0018). This suggests that while Paroxysmal AF represents a variable state of spatial instability, progression to the non-paroxysmal phenotype is characterized by a uniform “flattening” or homogenization of fluctuations dynamics across the chamber.

## DISCUSSION

Atrial fibrillation is a spatiotemporally complex electrical disorder of the heart, whose mechanisms remain intensely debated. Despite advances in high-density mapping, the mechanisms governing the persistence and termination of this disorder remain unresolved. A fundamental observation familiar to most electrophysiologists during AF recordings is the observation of bursts of organization and disorganization. Traditional signal processing approaches such as dominant frequency,(5) voltage(1) or entropy,(6) (5, 11–13), have the potential to smooth over nuanced characteristics like sudden periods of intermittent organised activity and non-stationary fluctuations. However, from the perspective of non-linear dynamics,(14) such fluctuations are not noise, these features may be important in understanding dynamical systems, like the human heart.

In this study, we aimed to systematically characterize the multifractal properties of AF electrograms for three key reasons: **(i)** to enable robust quantification of the fluctuating periods of organization and disorganization commonly observed during AF recordings in clinical procedures; **(ii)** to establish practical, reproducible metrics that capture these dynamic changes and could inform patient-specific treatment strategies; and **(iii)** to explore whether these multifractal signatures provide a mechanistic link to criticality, offering insight into how AF dynamics relate to phase transition phenomena within complex systems.

To achieve, this we developed analysis pipelines to allow analysis of a large prospectively registered, clinically-validated dataset of long recordings (>1 minute per site) of high-density mapping data in 106 patients.

The key findings of our study were:

i. **Multifractility:** Atrial fibrillation electrograms are multifractal in nature and can be described by broad scaling parameters.
ii. **Phenotype Differences:** Paroxysmal AF exhibits significantly higher multifractal fluctuations (c₂) compared to non-paroxysmal AF, indicating greater dynamical elasticity in early disease states.
iii. **Pharmacologic Modulation:** Flecainide selectively increases fluctuations in paroxysmal AF but has negligible effect in non-paroxysmal AF, highlighting phenotype-dependent responsiveness to based on levels of AF progression.
iv. **Termination Dynamics:** Immediately prior to spontaneous AF termination, fluctuations rises markedly, suggesting that elevated c₂ reflects proximity to a critical phase transition toward sinus rhythm.

### Loss of Fluctuations as a Signature of AF Progression

Our findings challenge the intuitive assumption that the progression of AF is driven by a simple increase in signal activation frequency or fractionation alone. Instead, we demonstrate that the transition from paroxysmal to non-paroxysmal AF is characterized by a paradoxical loss of multifractal fluctuations. While traditional metrics often describe AF progression as a move toward increasing disorganization, our application of the Wavelet Transform Modulus Maxima (WTMM) reveals that advanced disease states are more defined by a “dynamical smoothing” via reduction of the burst-like organized fluctuations (c_2_) that characterize conduction closer towards a more organised phase state. In this study, PAF exhibited a “rough” dynamical landscape, characterized by high values of fluctuations (c_2_) and significant spatial heterogeneity. In contrast, we observed that NPAF was associated with a global suppression of intermittent organization, and conduction undergoes dynamical homogenization. The spatial variogram analysis confirms the “texture” of the electrogram landscape flattens, suggesting disease state has created a uniformly disordered structure. In this state, the electrograms are less “bursty” and more monotonously turbulent.

This distinction is critical because it reframes how we view complexity in AF, where intermittent bursts of order and disorder (c_2_) represent a system attempting to transition back toward a macroscopically ordered state, and conversely, the loss of fluctuations in more advanced disease signals that the arrhythmia has settled into a stable, self-sustaining arrhythmia. Therefore, we propose that the loss of multifractal fluctuations (c_2_) serves as a novel biomarker able to discriminate between dynamics of state tending towards re-organisation (PAF pharmacological modulated AF and self-termination) and that of a rigid dynamical homogeneity of advanced disease.

### Multifractality as a property of heart homeostasis

To conceptualize why it is plausible for AF progression results in a loss of fluctuations, a possible interpretation of these findings is through the broader framework of multifractal criticality in the heart. The reduction of fluctuations in non-paroxysmal AF mirrors the loss of multifractal complexity observed in heart failure,(8, 15) or autonomic nervous system modulation,(16, 17) where healthy, heterogeneous heartbeat dynamics degrade into simpler, more monofractal patterns(8, 15–17). The multifractal complexity in heart rate observed in mammalian species is believed to occur as a consequence of self-organized criticality in heart rate modulation.(17)

The findings of this current study parallel these observations. Our data posits the fluctuations parameter (c_2_) acts as a metric for the system’s “distance from phase transition” where paroxysmal AF produces common intermittent bursts of organisation (relatively higher c_2_) suggest a system that potentially more proximate to a the phase transition boundary. In this critical state, the atria are not yet fully committed to the turbulent regime, but the tissue is in a constant state of fluctuation, rapidly switching between local organization and disorganization. The texture observed in the variograms of paroxysmal patients reflects this struggle, where the atria is a patchwork of electrograms complexity.

In contrast, the reduction of fluctuations in non-paroxysmal AF represents a system that has drifted further from the critical point. As disease progresses, atrial cell types, structures and connections remodel. This provides us with a conceptual paradox where in the advanced disease state, AF electrograms become more mathematically stable (lower variance in c_2_). However, this reflects disordered stability and homogeneity because disorder has become uniform everywhere. Advanced disease pushes the system away from the precipice of a phase transition, and it becomes locked into a stable attractor of high-dimensional turbulence, energetically distant from the conditions required for spontaneous reorganization.

### Pharmacological Intervention Shows Dynamical Elasticity in AF

To test the hypothesis that the loss of fluctuations represents a rigid, entrenched state, we used flecainide as a dynamical probe. By modulating (down) the excitability of the network, we were able systematically evaluate the systems close to the phase transition (paroxysmal) and those further into disease progression (non-paroxysmal).

The results reveal a striking bifurcation in response. In patients with paroxysmal AF, flecainide produced in a significant increase in conduction fluctuations (c2) consistent with known electrophysiologic effects of sodium channel blockade thus destabilizing singularities and forcing the system closer to the phase transition boundary. In contrast, the NPAF electrograms exhibited inelastic response to flecainide. Despite the same pharmacological intervention, our results show the multifractal spectrum remained unchanged after therapy. This lack of response suggests that the turbulent attractor is so deeply established that moderate changes in conduction velocity are insufficient for destabilization. In these patients, the fibrillatory network has become rigid and inelastic, and the turbulent dynamics are self-sustaining.

### Signal Fluctuations and the Transition to Order

An additional compelling observation lies in the dynamical signatures of spontaneous AF termination. Our analysis revealed that the spontaneous transition from fibrillation back to sinus rhythm is not an abrupt cessation of activity, but a structured dynamical event preceded by a significant spike in multifractal fluctuations (c_2_) and long-range organization (c_0_).

In statistical physics, this surge of fluctuations may represents a critical slowing down(18) as the complex system approaches a phase transition point at the bifurcation between the turbulent attractors of AF and the ordered attractors of sinus rhythm. At this phase transition point, the resilience of the AF state weakens and the system begins to experience macroscopic deviations as it explores the boundaries of its stability, until the correlation length increases exponentially and re-orders the system. The increase in c2 prior to spontaneous terminations provide further evidence for interpreting the dynamics of AF as a complex system that can be classified as being closer to, or further from termination boundary (criticality). A smoother and less intermittent signal reflects a system trapped deep within the turbulent regime, whereas high fluctuations represents an elastic system capable of engaging in self organisation to force a phase transition back to order.

### Reconciling Multifractality with Historical Mapping

Others have also observed these changes in dynamics prior to termination. Alcaraz et al employed non-linear methods (Sample entropy) to compare the dynamics of electrical activity immediately prior to termination to those of ongoing AF(19). Similar to our findings, they showed a reduction in entropy prior to termination and link this mechanistically to increase in **self-similarity in conduction.** With WTMM analysis, we can now extract additional information from these episodes to quantity the amount of self-similarity by evaluating the Holder exponents of the electrogram multifractal spectra.

The reduction of multifractal fluctuations (c2) as disease extends past criticality (paroxysmal - > non-paroxysmal) can be mechanistically reconciled with the well-established increase in Dominant Frequency (DF) described by Sanders et al. and Martins et al. Historical data posits that electrical remodelling causing downregulation of I_CaL_ and shortening of the effective refractory period accelerates vortex frequency, leading to higher DF(5, 20). However, this same remodelling process leads tissue to exhibit less vortex meandering and higher spatial stability. We propose that this stabilization across the atria manifests mathematically as a reduction in fluctuations due to vortex to fibrosis or other boundaries. While dominant frequency shows us that the rate of activation increases as disease progresses from criticality, the complexity of the variation decreases (Low c_2_). The system moves from a fragile, intermittent state (Paroxysmal) to a more robust, continuous state where the probability of spontaneous phase transition to sinus rhythm is minimized.

To conceptualize this complexity in AF, our group and others have modelled AF as scroll wave turbulence,(21) a state dynamically maintained by the creation and annihilation of topological defects known as phase singularities(22). Core to this theory is the a topological phase transition concept where the atria can pass between phases of ordered topology (sinus rhythm), bound vortices (atrial flutter) and finally AF representing the unbound vortex state, characterized by topological disorder and exponentially decaying spatial correlation(23). In this hypothesis, we propose that AF represents a complex system, and that as disease progresses, substrate remodelling pushes the system deeper into sustained, high-dimensional turbulence. Conversely, we also postulate that there must also exist a set of conditions when turbulence (ie AF) can transition back to ordered state (flutter, or sinus rhythm).

## Conclusion

Atrial Fibrillation is not a random process, but a complex dynamical system exhibiting robust multifractal scaling. We demonstrate that AF can be quantified with regard to closeness to the critical phase transition to sinus rhythm. We show that paroxysmal AF exists mathematically closer to the phase transition than non-paroxysmal AF, and verify this with observations from pharmacological intervention using flecainide, and evaluation of spontaneous termination events in a large clinical dataset of >1.4Million seconds of AF electrograms. We show that the ability for atrial tissues to exhibit both order and disorder can be used as a biomarker of closeness to the critical phase transition back to ordered state (sinus rhythm), and provide further validation of multifractal analysis methods for quantifying this.

## Abbreviations List

AF: Atrial Fibrillation
PAF: Paroxysmal Atrial Fibrillation
NPAF: Non-Paroxysmal Atrial Fibrillation
LA: Left Atrium / Left Atrial
RA: Right Atrium / Right Atrial
HD-Grid: High-Density Grid Catheter
WTMM: Wavelet Transform Modulus Maxima
ECG: Electrocardiogram / Electrocardiography
ERPs: Effective Refractory Periods (implied in discussion of remodelling)
DF: Dominant Frequency
CCDF: Complementary Cumulative Distribution Function
HREC: Human Research Ethics Committee
ANZCTR: Australia and New Zealand Clinical Trials Registry
ICaL: L-type Calcium Current
E(t): Burst-Energy Signal
SD: Standard Deviation
CI: Confidence Interval
IQR: Interquartile Range (if present in tables—appears implied)
NIH: National Institutes of Health (appears in context of grants)
Holder / Multifractal Parameters: 
c_₀_: Support Dimension
c_₁_: Spectrum Location
c_₂_: Fluctuations Parameter
τ(q): Scaling Function
f(α): Multifractal Spectrum

## REFERENCES

1. Rolf S, Dagres N, Hindricks G. Voltage-Based Ablation: The Growing Evidence for the Role of Individually Tailored Substrate Modification for Atrial Fibrillation. J Cardiovasc Electrophysiol. 2016;27(1):31–3.

2. Joglar José A, Chung Mina K, Armbruster Anastasia L, Benjamin Emelia J, Chyou Janice Y, Cronin Edmond M, et al. 2023 ACC/AHA/ACCP/HRS Guideline for the Diagnosis and Management of Atrial Fibrillation. JACC. 2024;83(1):109–279.

3. Van Gelder IC, Sanders P, Traykov V, Tzeis S, Kotecha D, Group ESCSD. 2024 ESC Guidelines for the management of atrial fibrillation. European Heart Journal. 2024;45(36):3314–414.

4. Reddy VY, Gerstenfeld EP, Natale A, Whang W, Cuoco FA, Patel C, et al. Pulsed Field or Conventional Thermal Ablation for Paroxysmal Atrial Fibrillation. New England Journal of Medicine. 2023;389(18):1660–71.

5. Sanders P, Berenfeld O, Hocini M, Jais P, Vaidyanathan R, Hsu LF, et al. Spectral analysis identifies sites of high-frequency activity maintaining atrial fibrillation in humans. Circulation. 2005;112(6):789–97.

6. Ganesan AN, Kuklik P, Lau DH, Brooks AG, Baumert M, Lim WW, et al. Bipolar electrogram Shannon entropy at sites of rotational activation: implications for ablation of atrial fibrillation. Circ Arrhythm Electrophysiol. 2013;6(1):48–57.

7. Zhao L, Li W, Yang C, Han J, Su Z, Zou Y. Multifractality and Network Analysis of Phase Transition. PLoS One. 2017;12(1):e0170467.

8. Ivanov PC, Amaral LAN, Goldberger AL, Havlin S, Rosenblum MG, Struzik ZR, et al. Multifractality in human heartbeat dynamics. Nature. 1999;399(6735):461–5.

9. Attuel G, Gerasimova-Chechkina E, Argoul F, Yahia H, Arneodo A. Multifractal Desynchronization of the Cardiac Excitable Cell Network During Atrial Fibrillation. I. Multifractal Analysis of Clinical Data. Front Physiol. 2017;8:1139.

10. Dharmaprani D, Jenkins E, Aguilar M, Quah JX, Lahiri A, Tiver K, et al. M/M/Infinity Birth-Death Processes – A Quantitative Representational Framework to Summarize and Explain Phase Singularity and Wavelet Dynamics in Atrial Fibrillation. Frontiers in Physiology. 2021;11:1786.

11. Ng J, Goldberger JJ. Understanding and interpreting dominant frequency analysis of AF electrograms. Journal of cardiovascular electrophysiology. 2007;18(6):680–5.

12. Nademanee K, McKenzie J, Kosar E, Schwab M, Sunsaneewitayakul B, Vasavakul T, et al. A new approach for catheter ablation of atrial fibrillation: mapping of the electrophysiologic substrate. J Am Coll Cardiol. 2004;43(11):2044–53.

13. Verma A, Jiang CY, Betts TR, Chen J, Deisenhofer I, Mantovan R, et al. Approaches to catheter ablation for persistent atrial fibrillation. The New England journal of medicine. 2015;372(19):1812–22.

14. Atienza F, Almendral J, Ormaetxe JM, Moya A, Martínez-Alday JD, Hernández-Madrid A, et al. Comparison of radiofrequency catheter ablation of drivers and circumferential pulmonary vein isolation in atrial fibrillation: a noninferiority randomized multicenter RADAR-AF trial. (1558-3597 (Electronic)).

15. Goldberger AL, Amaral LAN, Hausdorff JM, Ivanov PC, Peng CK, Stanley HE. Fractal dynamics in physiology: alterations with disease and aging. Proceedings of the national academy of sciences. 2002;99(suppl_1):2466–72.

16. Moghtadaei M, Dorey TW, Rose RA. Evaluation of non-linear heart rate variability using multi-scale multi-fractal detrended fluctuation analysis in mice: Roles of the autonomic nervous system and sinoatrial node. Front Physiol. 2022;13:970393.

17. Tagirova Sirenko S, Tsutsui K, Tarasov KV, Yang D, Wirth AN, Maltsev VA, et al. Self-Similar Synchronization of Calcium and Membrane Potential Transitions During Action Potential Cycles Predict Heart Rate Across Species. JACC Clin Electrophysiol. 2021;7(11):1331–44.

18. Maturana MI, Meisel C, Dell K, Karoly PJ, D’Souza W, Grayden DB, et al. Critical slowing down as a biomarker for seizure susceptibility. Nature Communications. 2020;11(1):2172.

19. Alcaraz R, Rieta JJ. A novel application of sample entropy to the electrocardiogram of atrial fibrillation. Nonlinear Analysis: Real World Applications. 2010;11(2):1026–35.

20. Martins RP, Kaur K, Hwang E, Ramirez RJ, Willis BC, Filgueiras-Rama D, et al. Dominant frequency increase rate predicts transition from paroxysmal to long-term persistent atrial fibrillation. Circulation. 2014;129(14):1472–82.

21. Christoph J, Chebbok M, Richter C, Schröder-Schetelig J, Bittihn P, Stein S, et al. Electromechanical vortex filaments during cardiac fibrillation. Nature. 2018;555(7698):667.

22. Dharmaprani D, Tiver K, Salari Shahrbabaki S, Jenkins EV, Chapman D, Strong C, et al. Observable Atrial and Ventricular Fibrillation Episode Durations Are Conformant With a Power Law Based on System Size and Spatial Synchronization. Circ Arrhythm Electrophysiol. 2024;17(7):e012684.

23. Ganesan AN, Kuklik P, Nattel S. A topological hypothesis for atrial fibrilllation, atrial flutter and focal atrial tachycardia: comparison and contrast with Kosterlitz-Thouless physics. Frontiers in Network Physiology. 2026;Volume 5 - 2025.

